# VGLUT2 functions as a differential marker for hippocampal output neurons

**DOI:** 10.1101/348227

**Authors:** Christian Wozny, Prateep Beed, Noam Nitzan, Yona Pössnecker, Benjamin R. Rost, Dietmar Schmitz

## Abstract

The subiculum is the gatekeeper between the hippocampus and cortical areas. Yet, the lack of a pyramidal cell-specific marker gene has made the analysis of the subicular area very difficult. Here we report that the vesicular-glutamate transporter 2 (VGLUT2) functions as a specific marker gene for subicular burst-firing neurons, and demonstrate that VGLUT2-Cre mice allow for Channelrhodopsin-2 (ChR2)-assisted connectivity analysis.

## Main text

The hippocampal formation consists of anatomically defined brain areas such as the dentate gyrus (DG), the cornu ammonis (area CA1-CA3 in rodents), the subiculum (SUB), pre- and parasubiculum, and the entorhinal cortices (EC), all of which fulfill a variety of different tasks. Well documented is the role of these hippocampal subregions in spatial navigation.

The SUB is the main hippocampal output structure and functions as a relay between the hippocampus proper and the EC. Two different types of subicular pyramidal neurons have been described based on their intrinsic firing pattern as burst- or regular-firing neurons ^1–3^. These neurons project to different brain regions: burst-firing neurons to the medial EC, the presubiculum, the retrosplenial cortex, and to the hypothalamus, whereas mostly regular-firing neurons project to the amygdala, the lateral EC and the nucleus accumbens ^4^. Remarkably, CA1 inputs to these two cell-types express different forms of synaptic plasticity ^5–7^. The intrinsic firing pattern, however, is not static, but might be modulated by neuronal activity ^8^. Until now, the lack of cell-specific marker genes for burst- and regular-firing neurons has, however, hampered the application of state-of-art circuit analysis tools such as Channelrhodopsin-2 (ChR2), which would allow to redefine the cortico-hippocampal wiring diagram and to further disentangle the role of these particular neurons.

To identify promising candidate genes for SUB-specific expression pattern we screened public repositories such as the Allen Brain Atlas. A differential search between area CA1 of the hippocampus and the SUB further aided to narrow the number of genes, and revealed VGLUT2 as one of the most promising candidates (Supplementary Fig. 1).

Injection of an adeno-associated virus (AAV) encoding Cre-dependent DIO (double-floxed inverted orientation) ChR2(H134R)-mCherry into the SUB resulted in localized expression of ChR2-mCherry (Fig. 1a-c). To test the functional expression of ChR2 we recorded light-evoked responses in the presence of synaptic blockers (Fig. 1d,e).

**Figure 1:**
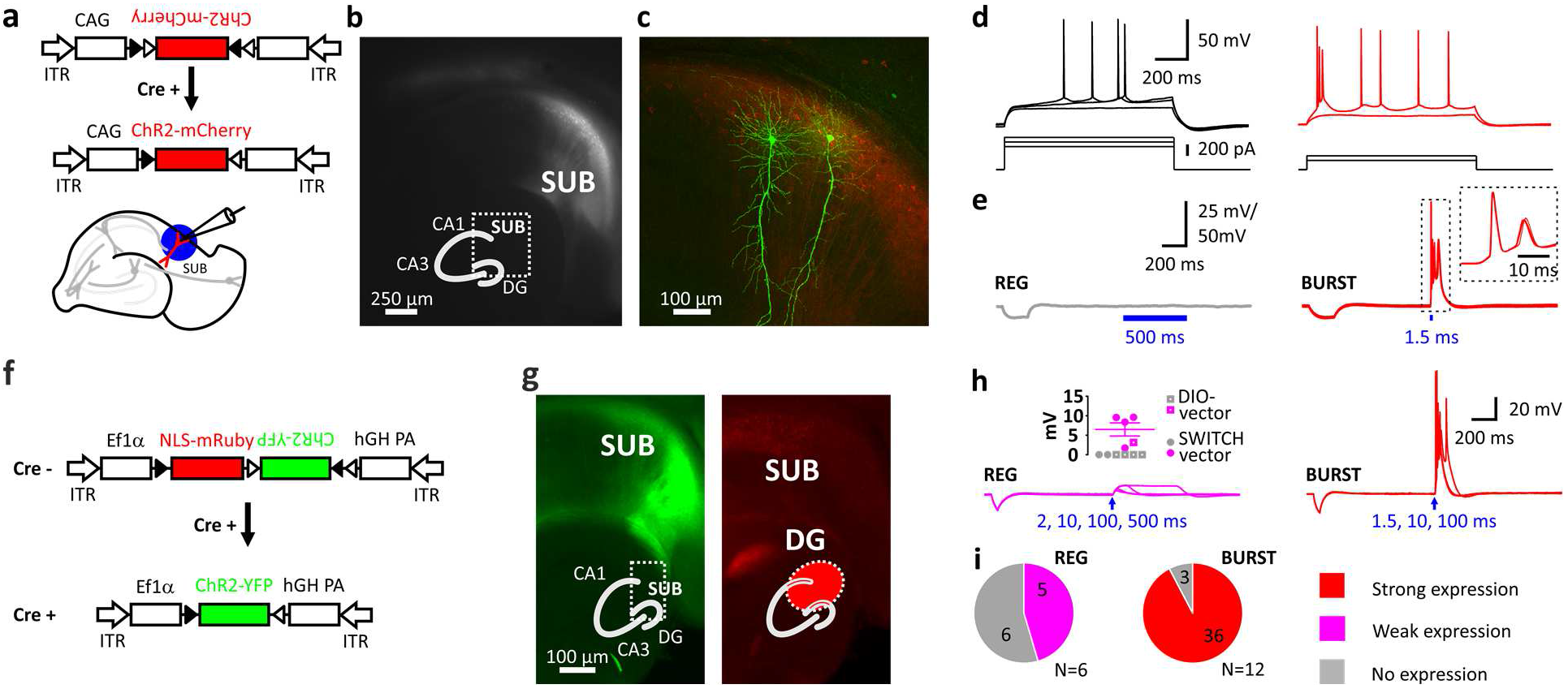
Selective expression of Cre-dependent ChR2 in subicular burst-firing neurons in VGLUT2-Cre mice. **a,** Injection of AAVs into the subiculum (SUB) of VGLUT2-Cre mice. Schematic showing the Cre-conditional DIO-ChR2-mCherry construct and the slice preparation. Blue circle illustrates blue light illumination. **b,** Live image of mCherry fluorescence showing mCherry-positive neurons in the SUB. Inset shows schematic of hippocampus with DG, CA3, CA1 and SUB. **c,** Maximum projection of confocal z-stacks of biocytin-filled neurons and virally induced expression of the red fluorophore. **d,** Firing pattern of a subicular regular-firing (REG, left), and of a subicular burst-firing neuron (BURST, right). **e,** Left: Average of 10 traces using a 500ms light pulse demonstrated no expression of ChR2 in REG neuron. Right: In contrast, a 1.5ms light pulse elicited action-potential firing in subicular BURST neuron; inset shows 10 consecutive traces. Recordings were performed in the presence of blockers of synaptic transmission. **f,** Switch vector design allowing NLS-mRuby expression in Cre-negative and ChR2-YFP expression in Cre-positive neurons. **g,** Left: EYFP signal is restricted to SUB, whereas NLS-mRuby is seen in the distal CA1 region, SUB and the DG (on the right). Infected area indicated in red in inset. **h,** Left: Blue-light stimulation of up to 500 ms (indicated by arrow) elicited no or small depolarizations of < 10 mV due to no or weak expression of ChR2 in regular-firing (REG) neurons. Inset shows evoked depolarization (n=5; N=3 mice). Closed circles indicate mice infected with Switch vector, color code as in **i.** Right: Strong expression of ChR2 resulted in action-potential firing in BURST neurons (n=36 out of 39 neurons; N=12 mice). **i,** None of the REG neurons showed strong expression of ChR2, which means none fired APs (n=11 neurons; N=6 mice), in contrast to 36 out of 39 BURST neurons showing light-induced AP-firing (N=12 mice). Data pooled from both Cre-conditional constructs.

We first classified SUB neurons in respect to their intrinsic firing pattern into burst- and regular firing neurons (Fig. 1c). Expression of ChR2 nicely coincided with the intrinsic firing pattern as subsequent light stimulation elicited action potentials (AP) only in intrinsically burst-firing neurons (Fig. 1e,h,i). Post-hoc confocal imaging of the biocytin-labeled recorded neurons confirmed the typical characteristics of pyramidal neurons (Fig. 1c), as well as co-localization of the biocytin signal and the virally induced expression of ChR2-mCherry (Supplementary Fig. S2).

We next created a Cre-dependent Switch vector that allowed us to identify both Cre-positive neurons by ChR2-YFP expression, as well as Cre-negative neurons by a red fluorophore (Fig. 1f). In Cre-negative cells, the Ef1α promoter drives expression of mRuby2 fused to a nuclear-localization sequence (NLS), whereas the ChR2-YFP coding sequence is reversely orientated.

Cre-mediated recombination of the construct removes the NLS-mRuby2 coding sequence and initiates ChR2-YFP expression by inverting the ChR2-YFP sequence. This technique allowed us to identify the infected brain area more precisely, and also to distinguish infected from uninfected neurons (Fig. 1g).

Regarding the expression of ChR2 in subicular neurons the response patterns upon blue light illumination were further analyzed and grouped into three categories: (i) strong, (ii) weak and (iii) no response. A strong response was considered to be an AP. In 36 out of 39 burst-firing neurons light elicited an AP (AP size: 108.7 ± 1.8 mV, n = 36; Fig. 1h). A weak response was smaller than 10 mV following either a 10, 100, 200 or 500 ms light pulse. Five out of eleven regular-firing neurons showed such a response (6.5 ± 1.7 mV, range 1.7 – 9.6 mV, n = 5; see inset Fig. 1h). No expression of ChR2, and therefore no response to light was evident in 3 out 39 burst-firing (8%), and in 6 out of 11 regular-firing neurons (55%; Fig. 1i).

Previously, subicular neurons have been shown to project to the deep layers of the entorhinal cortex ^9^. More recently the wiring has been refined demonstrating that subicular (and CA1) pyramidal neurons project onto Coup-TF interacting protein 2 (CTIP2)-expressing neurons located in layer 5b of the EC ^10^. However, whether these subicular neurons express VGLUT2 is unclear.

**Figure 2:**
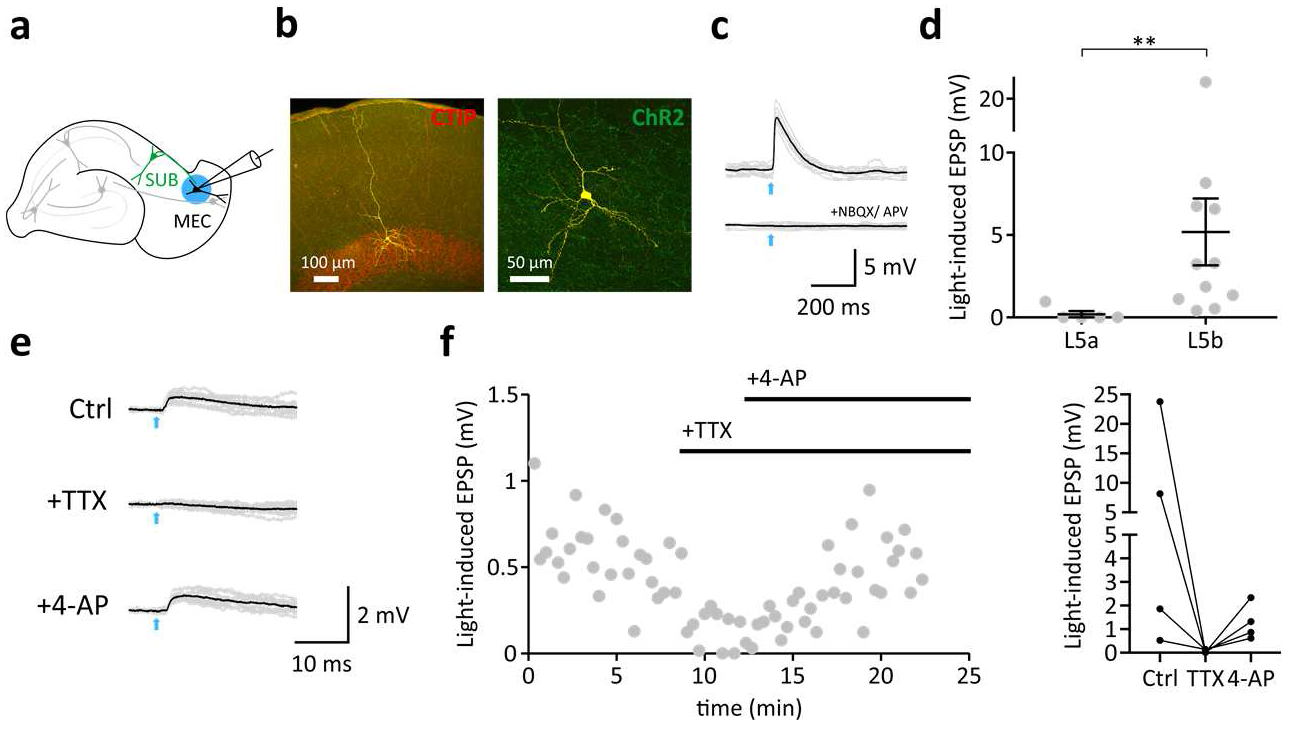
Monosynaptic connections from the SUB to mEC L5b neurons. **a,** Schematic showing ChR2-infected burst-firing neuron in the SUB, and subsequent whole-cell recordings from mEC neuron. **b,** Anatomical reconstruction of an EC L5b pyramidal neuron. Yellow: Biocytin-streptavidin-Alexa647 signal. Red: Immunoreactivity against CTIP. Green: ChR2-YFP. **c,d,** Light-induced postsynaptic responses (EPSPs; illumination indicated by an arrow, 5ms) of mEC L5a (n=5, N=4 mice) and L5b pyramidal neurons (**d,** n=11, N=6 mice, Mann-Whitney test, p<0.01). **e,** Light stimulation (indicated by an arrow, 5 ms) elicited an EPSP (left), which was blocked by TTX, and rescued with 4-AP confirming monosynaptic nature of the signal. **f,** Left, time course. Right, individual 4-AP experiments (n = 4, N = 3 mice).

We therefore recorded from layer 5b neurons in the EC and applied ChR2-assisted mapping circuit in VGLUT2-Cre mice to monitor efferent monosynaptic connections of subicular burst-firing neurons (Fig. 2-d). Subicular efferents were stimulated using blue light pulses (2-5 ms; Fig. 2c). Application of either glutamate receptor blockers (Fig. 2c), or the sodium channel blocker tetrodotoxin (TTX; Fig. 2e,f) abolished the response, confirming that the synaptic response was driven by AP-triggered transmitter release. Addition of the potassium channel blocker 4-aminopyridine (4-AP) rescued the light-evoked EPSP (Fig. 2e,f), proving the monosynaptic nature of the synaptic input. In contrast, and consistent with previous reports ^10^, layer 5A neurons did not receive inputs from subicular burst-firing neurons (Figure 2d; Mann-Whitney test p<0.01).

Here we report the identification of VGLUT2 as a marker of subicular burst-firing neurons. By utilizing VGLUT2-Cre mice in combination with viral gene delivery strategies of Cre-dependent ChR2-expression constructs we demonstrate that VGLUT2 expression is mainly found in subicular burst-firing neurons. This versatile tool allows microcircuit analysis confirming and extending previous results of where hippocampal output neurons synapse onto mEC L5b neurons ^9–12^.

Recently, the restricted expression of fibronectin-1 (FN1) in the dorsal subiculum was utilized to generate a mouse line expressing Cre specifically in dorsal SUB neurons. Subsequently, optogenetic manipulations were performed to address the role of the dorsal SUB in memory formation ^13^. Another recent study divided proximal and distal subicular pyramidal neurons ^14^. Proximal subicular neurons express neuronatin (Nnat), whereas neurotensin (Nts) is found in distal neurons. However, it is currently not known whether these marker genes specifically label subpopulations of subicular principal neurons

Földy et al. performed a single-cell transcriptome study of subicular pyramidal neurons ^15^. Following electrophysiological characterization the mRNA of burst- and regular-firing neurons was collected, and sequenced. There was no difference in the number of detected genes between these two types of subicular neurons, and only a small number of exclusively expressed genes were found, however, none of these were further validated as a functional genetic marker. We used a differential approach. Viruses were used to infect Cre- and non-Cre-recombinase expressing neurons. In our hands, over 90% of the recorded burst-firing neurons expressed high amounts of ChR2, whereas none of the subicular regular-firing neurons expressed sufficient amounts of ChR2 to drive AP firing following blue light illumination with varying lengths.

Of note, the expression of VGLUT1 and VGLUT2 in the brain is thought to be complementary: VGLUT1 is mainly expressed in cortical areas, whereas VGLUT2 is expressed in subcortical areas such as the thalamus, amygdala or hypothalamus ^16–18^. A few brain areas including the SUB, however, seem to express both, VGLUT1 and VGLUT2 ^19, 20^. Whether VGLUT1-positive neurons in the SUB are mainly of the regular-firing type has to be determined, as has the role of both types of subicular neurons during behavior.

## Acknowledgements

We would like to thank Susanne Rieckmann and Anke Schönherr for excellent technical assistance.

## Notes

This work was supported by a Wellcome Trust Seed Award to CW (205917/z/17/Z), by the NeuroCure Clusters of Excellence, the DZNE, the Einstein Foundation, and the Deutsche Forschungsgemeinschaft to DS (EXC257, SFB858, SPP1665), by the SPP1926 (‘Next Generation Optogenetics’) to BRR, and by the Stiftung Charité to PB.

